# A hand-off of DNA between archaeal polymerases allows high-fidelity replication to resume at a discrete intermediate three bases past 8-oxoguanine

**DOI:** 10.1101/2020.06.17.157875

**Authors:** Matthew T. Cranford, Joseph D. Kaszubowski, Michael A. Trakselis

## Abstract

During DNA replication, the presence of 8-oxoguanine (8-oxoG) lesions in the template strand cause the high-fidelity (HiFi) DNA polymerase (Pol) to stall. An early response to 8-oxoG lesions involves ‘on-the-fly’ translesion synthesis (TLS), in which a specialized TLS Pol is recruited and replaces the stalled HiFi Pol for bypass of the lesion. The length of TLS must be long enough for effective bypass, but it must also be regulated to minimize replication errors by the TLS Pol. The exact position where the TLS Pol ends and the HiFi Pol resumes (*i.e*. the length of the TLS patch) has not been described. We use steady-state and pre-steady-state kinetic assays to characterize lesion bypass intermediates formed by different archaeal polymerase holoenzyme complexes that include PCNA123 and RFC. After bypass of 8-oxoG by TLS PolY, products accumulate at the template position three base pairs beyond the lesion. PolY is catalytically poor for subsequent extension from this +3 position beyond 8-oxoG, but this inefficiency is overcome by rapid extension of HiFi PolB1. The reciprocation of Pol activities at this intermediate indicates a defined position where TLS Pol extension is limited and where the DNA substrate is handed back to the HiFi Pol after bypass of 8-oxoG.

## Introduction

Accurate replication of genomic DNA is an elaborate cellular process which is essential to the propagation of life. The double helix is first unwound to form a replication fork where the two strands are primed for replication. A high-fidelity (HiFi) DNA polymerase (Pol) then associates with a processivity clamp to form a replicative holoenzyme (HE) complex which utilizes the single strands of the DNA substrate as a template for accurate duplication of the genome. At times, the molecular information of the template DNA is exposed to damaging agents and becomes chemically modified to form a variety of DNA lesions. DNA repair pathways (including nucleotide-excision repair and base-excision repair) allow for correction of these genomic lesions, yet, excess exposure to DNA damaging agents allows lesions to persist and then be encountered during replication. These template lesions cause HiFi Pols to stall, leading to genomic instability.

However, a specialized set of evolutionarily conserved Y-family Pols can perform translesion synthesis (TLS) by incorporating deoxynucleotides (dNTPs) across template lesions (1). Although these TLS Pols are prone to errors during replication (2), they provide a pathway for DNA damage tolerance and allow replication to proceed in the presence of unrepaired DNA lesions. Previous studies have implicated two DNA damage tolerance pathways which involve TLS Pols: by ‘re-priming,’ a new primer is synthesized downstream of the replication-stalling lesion (3-5). This allows the HiFi Pol to skip the lesion and continue synthesis downstream. However, this discontinuous process creates a single-stranded gap which must be resolved by either template-switching homology-directed gap repair (a TLS-independent pathway) or post-replicative gap filling by TLS Pols (6). Alternatively, TLS may occur ‘on-the-fly,’ in which a TLS Pol is recruited directly to a stalled replication fork to perform lesion bypass prior to a re-priming event (7). Recent *in vitro* studies have suggested that this continuous ‘on-the-fly’ TLS pathway is an early response to replication stalling lesions, whereas re-priming serves as a slower mechanism which is likely utilized during conditions of excessive DNA damage (8-10).

The model for ‘on-the-fly’ TLS presents two distinct events where the DNA substrate is ‘handed-off’ between the HiFi and TLS Pols. Upon encountering a lesion, a HiFi Pol stalls one template base prior to the lesion (−1) as a result of competing polymerase and exonuclease activity (11). From this stalled position, the DNA must be handed-off from the stalled HiFi Pol to a TLS Pol, allowing for bypass of the template lesion. The mechanism of this hand-off may occur indirectly by distributive ‘exchange,’ where an active Pol fully dissociates from the DNA prior to equilibrium-driven recruitment of the next Pol from solution (12); or, the DNA may be passed directly through a ‘switch’ within a coupled supraholoenzyme (SHE) complex consisting of multiple Pols bound to a single processivity clamp (*i.e*. the tool belt model) (13-16). The TLS Pol synthesizes a nascent segment of DNA which bypasses the lesion and extends to an intermediate position downstream. However, the exact length of this TLS patch and the position of the second hand-off (to resume synthesis by the HiFi Pol) has not been investigated. It is understood that the length of the TLS patch must be long enough to prevent lesion recognition and exonucleolytic degradation by the HiFi Pol, but continued extension beyond the lesion by low-fidelity TLS Pols must be limited to minimize error-prone synthesis (17). Limitations to TLS Pol extension may indeed be explained by their characteristically low processivity (18), but the complex multieuilibria of the cell provides multiple factors which may regulate the TLS Pol. A previous study demonstrated that a TLS patch length of 5 nucleotides beyond an acetylaminofluorine adduct was sufficient for HiFi *E. coli* Pol III to resume replication; however this is not implicated as the position of the second hand-off, as TLS Pol V is capable of much longer processive stretches of synthesis (19). In eukaryotes, synthesis of this TLS patch is more intricate, involving posttranslational modifications of PCNA (20) and additional hand-offs among the various TLS inserter and extender Pols which possess unique activities for inserting across and extending beyond different types of DNA lesions, respectively (21). This is further complicated by a recent study revealing competition between Pold and Polh in a stalled replisome that is decoupled from the CMG, even on a leading-strand lesion (10). Nevertheless, the extent of combined TLS Pol activity beyond the lesion is not fully understood. This current knowledge gap encourages additional investigation into the molecular mechanisms which limit error-prone TLS extension after lesion bypass and restore accurate HiFi replication.

PolY (also referred to as Dpo4) acts as the TLS Pol in the archaeon *Saccharolobus solfataricus* (*Sso*) (22). This model enzyme has been well studied structurally (23-26) and kinetically (24,27,28), and its legacy has provided a basis for our current understanding of the various eukaryotic TLS Pols (29). PolY adopts the canonical ‘right-handed’ Pol structure with ‘Fingers,’ ‘Palm,’ and ‘Thumb’ subdomains (23,30). As a Y-family polymerase, PolY also contains the unique Little-Finger (LF) domain which plays a central role in TLS activity (23,31). In response to 8-oxoguanine (8-oxoG) lesions, PolY is remarkably efficient and accurate for non-mutagenic insertion of dCTP (24,28). Other Pols are prone to incorrect insertion of dATP across 8-oxoG by formation of a Hoogsteen base pair, resulting in G-to-T transversion mutations (32). However, R332 (in the LF of PolY) stabilizes the *anti*-conformation of 8-oxoG in the PolY active site, allowing accurate Watson-Crick base pairing and efficient insertion of dCTP (24,25,33).

Downstream of 8-oxoG, PolY is capable of extension activity (24,28). However, the extent of PolY’s catalytic role beyond the lesion in the presence of other replisome components has not been investigated fully. This prompts examination into how these ‘right-handed’ Pols ‘hand-off’ the DNA in response to lesions. One previous kinetic study included PolB1 (the HiFi Pol in *Sso*) to evaluate its role in extension after PolY bypass of 8-oxoG (28), albeit in the absence of the heterotrimeric processivity clamp PCNA123 (34,35) and clamp loader complex RFC (36,37). PolB1 polymerase and exonuclease activities were unperturbed by the template 8-oxoG lesion on a DNA substrate with a primer terminus that was extended 1 base pair downstream (+1) of the lesion. This +1 position is highlighted as a kinetically favorable location for the second Pol hand-off. However, the authors ultimately conclude that inclusion of PCNA123 may alter the balance of bypass and extension activities by PolY and PolB1, and this mediation may dictate the position where the DNA is handed off between Pols after bypass of 8-oxoG (28).

Building upon this kinetic framework, we have determined that PolB1 hands-off the template to PolY to perform lesion bypass across 8-oxoG and extends to a position three base pairs beyond the lesion (+3). Additional extension by PolY from this position is inefficient, leading to accumulation of a +3 intermediate product. Within a SHE complex (composed of PolB1, PolY and PCNA123), accumulation of the +3 intermediate is reduced as a result of a second hand-off to resume rapid extension activity of PolB1. Collectively, this identifies a specific position for the second Pol hand-off where HiFi replication resumes after TLS bypass of an 8-oxoG lesion during ‘on-the-fly’ TLS.

## Results

### PolY and PolB1 have complementing activities for bypass of and extension from an 8-oxoG lesion

To measure lesion bypass activity, different Pol HE complexes were assembled by pre-loading PolB1 and/or PolY with PCNA123 on a damaged DNA substrate containing an 8-oxoG lesion. Primers of progressing length were utilized in order to monitor lesion bypass or extension activities from different starting positions relative to the site of the template lesion (**Fig. 1A**). Reactions were initiated upon addition of dNTPs, Mg^2+^, and an excess amount of non-specific salmon sperm DNA (spDNA) as a trap to alter the multiequilibria of polymerase ‘exchange.’ As a pre-trapping control, spDNA was pre-incubated with the DNA substrate prior to reaction initiation (**Supplemental Fig. S1**). At 1 mg/ml of spDNA, pre-trapped Pol complexes are not completely trapped in solution, suggesting that this concentration of spDNA does not abolish but slows the rate of distributive Pol exchange from solution. On the running start primer p18 (−5), synthesis takes place prior to encountering the 8-oxoG lesion. However, to compare lesion bypass activity, only the signal of ‘Bypassed’ products beyond the position of the lesion were quantified (**Fig. 1B**, brackets). From the running start position, the PolB1 holoenzyme (B1HE; lanes 1-3) is completely stalled at the 8-oxoG lesion with negligible signal beyond the lesion (purple). As expected, reactions with the PolY holoenzyme (YHE, lanes 4-6) demonstrate lesion bypass activity with 39.8 ± 1.5% of products extended beyond the lesion (red). Upon examining the gel image, the distribution of lesion bypass products by the YHE appear to be enriched at an intermediate position downstream of the lesion (**Fig. 1B**, arrow). This indicates that PolY is capable of TLS, but it exhibits poor processivity and limited extension beyond the 8-oxoG lesion. Lastly, formation of the SHE complex, containing both PolB1 and PolY (lanes 7-9), yields significantly more (52.6 ± 3.3 %; blue) lesion bypass products compared to the YHE. Additionally, the intermediate observed in the YHE reactions is not apparent in the lesion bypass products for the SHE, and more products are elongated to FL.

**Fig. 1.**
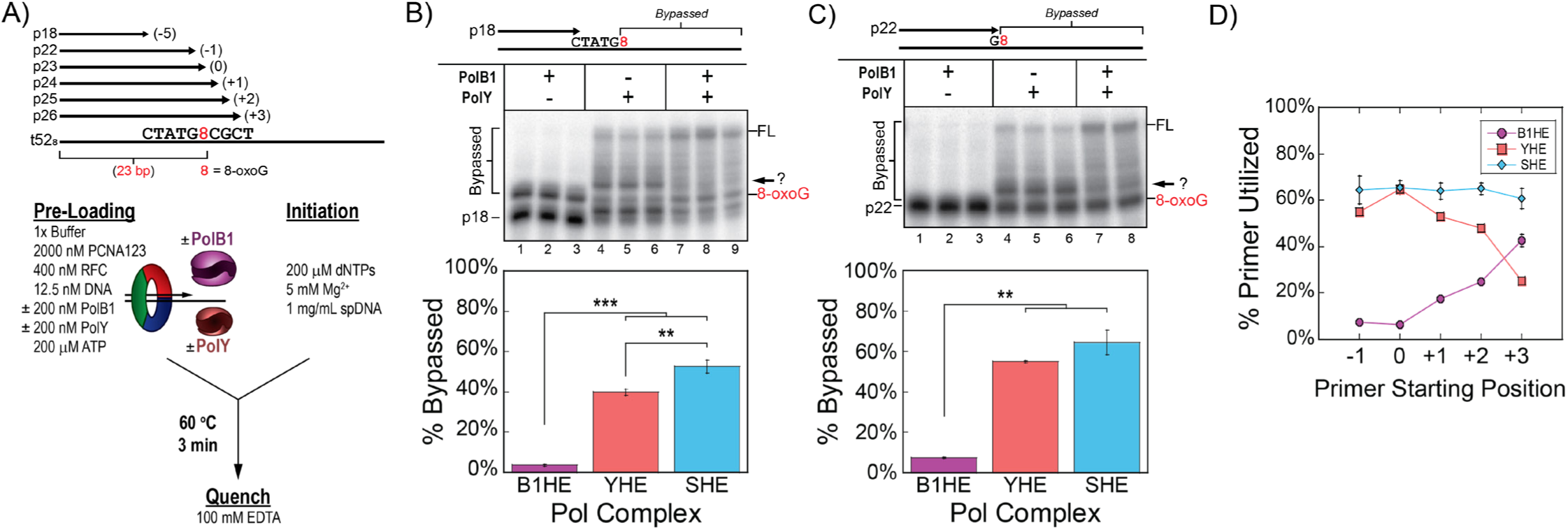
PolY and PolB1 have complementing activities for bypass and extension from an 8-oxoG lesion. A) A 52-mer DNA template containing an 8-oxoG lesion at position 23 (from the 3’ end) was annealed to primers of progressing lengths. DNA sequences for each oligo are listed in **Supplemental Table 1**. Steady-state lesion bypass assays were performed by pre-loading the indicated DNA substrate with different replication subassemblies at indicated (final) concentrations (see Materials and Methods). Lesion bypass reactions performed on B) the running start substrate and C) the stalled substrates were resolved by low-resolution denaturing PAGE. Products formed by the indicated Pol complex were quantified for percentage of ‘Bypassed’ products beyond the lesion position. D) Steady-state extension assays were performed on substrates with longer starting primers. Products quantified for the percentage of primer that was utilized by each Pol complex (B1HE, purple -•-; YHE, red -■-; SHE, blue -♦-). Gel images are in **Supplemental Fig. S2**. FL indicates the position of full-length product, and arrows indicate an intermediate product downstream of 8-oxoG. Error bars represent the standard deviation from three independent replicates.

We repeated this lesion bypass assay on a DNA substrate simulating a stalled position using primer p22 (−1; **Fig. 1C**). Again, the B1HE (lanes 1-3) is stalled by 8-oxoG (purple), while the YHE (lanes 4-6) is capable of lesion bypass (55.0 ± 0.5 %, red). As observed in **Fig. 1B**, most of the lesion bypass products by the YHE are concentrated at a position beyond the lesion (**Fig. 1C**, arrow). Finally, though the SHE (lanes 7-8) does not yield a significant increase in total bypassed products (64.3 ± 6.0 %, blue) relative to the YHE, most of the bypassed products are extended to the end of the substrate and the observed intermediate disappears. Thus, in response to 8-oxoG, bypass and extension from the lesion are most efficient when both PolB1 and PolY are present.

We continued these lesion bypass assays for each Pol complex using DNA substrates with progressively longer primers and quantified the percentage of primer that was utilized for each substrate (*i.e*. extended by at least one base pair; **Fig. 1D**, gel images in **Supplemental Fig. S2A-D**). In this way, we can compare the activities of each Pol complex on substrates with different starting positions relative to the 8-oxoG lesion. Activity of the B1HE is negligible on primers p22 and p23, which start before (−1) or at (0) the site of the lesion. However, activity of the B1HE gradually increases on primers further downstream of the lesion (purple circles). Remarkably, products from the YHE exhibit the opposite trend; activity is elevated on primers before or at the lesion and gradually decreases on primers which require extension from the lesion (red squares). Finally, activity of the SHE consistently yields about 65% of extension on all primers (blue diamonds). Unlike reactions with individual Pol HEs, the magnitude of primer extension by the SHE complex is independent of starting primer length.

The apparent inverse in activities relative to the 8-oxoG position by the YHE and B1HE highlights the complementing roles for PolY and PolB1. This is further supported by the SHE reactions, where the combined Pol activities yield increased levels of bypass and extension. These data also point to a specific position where the substrate is handed off between the Pols in the SHE reactions. The reciprocation of Pol activities occurs on a primer terminus positioned 3 base pairs downstream of the lesion. This leads us to hypothesize that the +3 position is the site of the second hand-off, from TLS PolY to HiFi PolB1, after bypass of an 8-oxoG lesion.

### Lesion bypass by PolY yields a major intermediate 3 base pairs downstream of the 8-oxoG lesion

In order to visualize discrete lesion bypass intermediates, representative products from the previous reactions were resolved by high-resolution denaturing PAGE (**Fig. 2A**). This gel image depicts the same activity trends summarized in **Figure 1D**, but it also grants a more precise examination of lesion bypass and extension products formed by each Pol complex. Notably, lesion bypass products by the YHE are specifically enriched at the +3 position, regardless of the primer starting position (**Fig. 2A**, arrow). The consistent position of this enriched YHE intermediate indicates that it is not a product of an ordinary processive step by PolY. If so, the position of the enriched intermediate would shift along the substrate in relation to the primer starting length. This observation indicates a specific perturbation in polymerase activity of PolY when positioned 3 base pairs downstream of an 8-oxoG lesion.

**Fig. 2.**
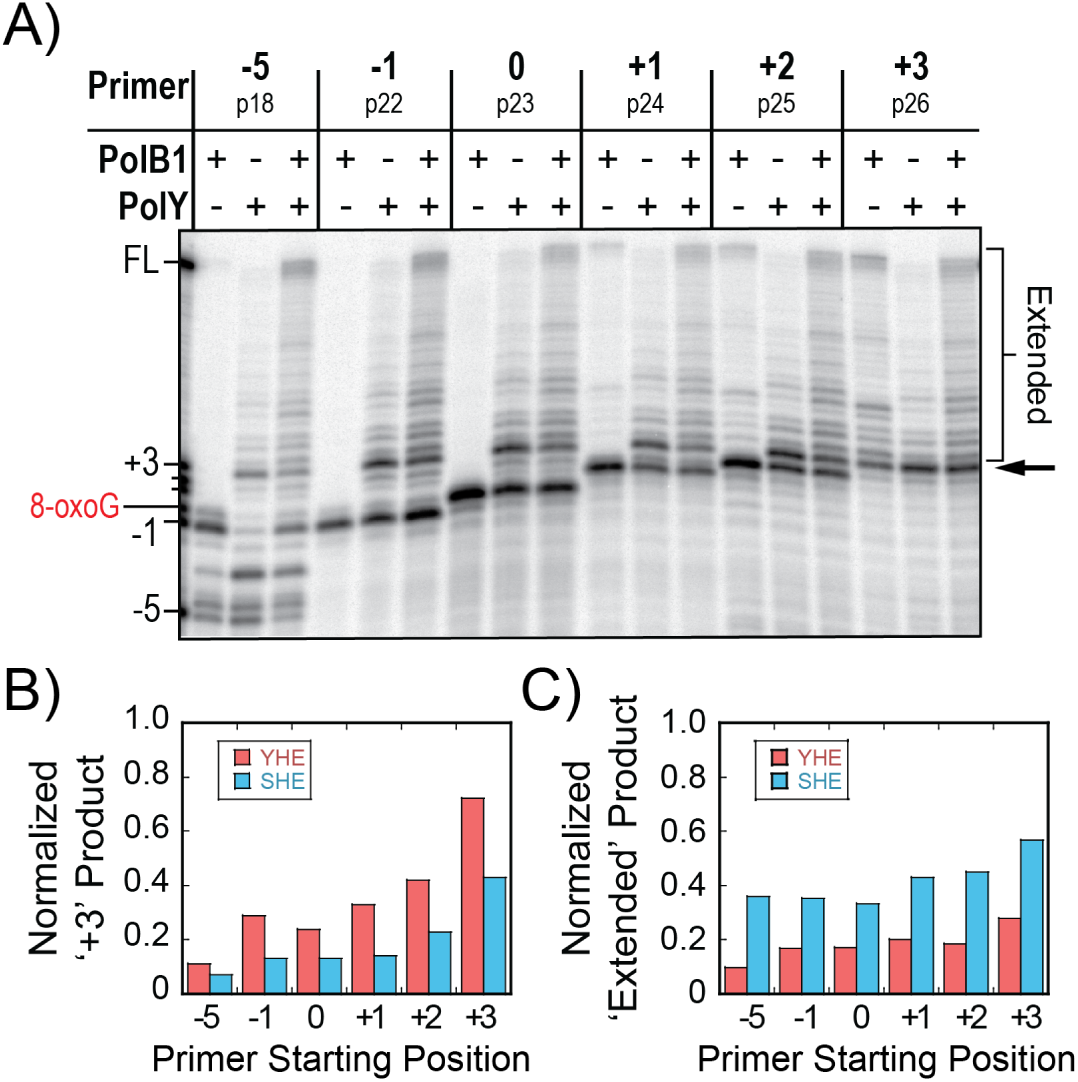
Lesion bypass by PolY yields a major intermediate 3 base pairs downstream of the 8-oxoG lesion. A) Representative lesion bypass and extension products by different Pol complexes (from **Fig. 1** and **Supplemental Fig. S2**) were resolved by high-resolution denaturing PAGE. The arrow indicates the intermediate product 3 base pairs beyond (+3) the 8-oxoG lesion, and ‘Extended’ indicates all products beyond the +3 position. The normalized fraction of B) ‘+3’ product bands and C) ‘Extended’ products for the YHE (red) or SHE (blue) were plotted from indicated primer starting positions.

As previously noted, lesion bypass with the SHE yields an apparent decrease in the intermediate observed in bypass by the YHE complexes and an apparent increase in products extended to FL. In order to compare the relative amount of +3 intermediate formed on each substrate, the signal of the +3 intermediate product bands (**Fig. 2A**, arrow) from the YHE and SHE reactions were normalized to their respective lanes (**Fig. 2B**). When compared, the magnitude of the +3 intermediate signals from the SHE reactions (blue) is reduced by approximately half relative to the YHE reactions (red) on corresponding DNA substrates. Additionally, upon comparing the normalized fraction of extended products (**Fig. 2A**, bracket), the SHE reactions (blue) yield about twice as much product that is extended beyond the +3 position relative to the YHE (red; **Fig. 2C**). Altogether, these data indicate that PolY conducts bypass of an 8-oxoG lesion and is disrupted at a position 3 base pairs downstream, and the presence of PolB1 promotes extension from this intermediate position.

### Kinetics of lesion bypass intermediates demonstrate reduced accumulation of the +3 intermediate upon formation of the SHE

After recognizing the +3 position as a major intermediate for PolY, we examined the direct kinetics of the intermediate without altering the equilibria by spDNA. In order to acquire faster time points, reactions were operated through an RQF-3 (KinTek) rapid quench instrument at 40 °C. PolY was introduced to the reaction by either syringe A or B (**Fig. 3A**, *Schemes i* or *ii*). By changing the position where PolY is introduced to the reaction, we can evaluate how the observed rate of lesion bypass changes under different pre-loading conditions. Any products which extend from this primer were quantified as ‘Total Bypass’ (**Fig. 3B**, left bracket), as the 8-oxoG lesion is the first templating base from the stalled primer p22 (−1) terminus. Further, all extension products were classified into three categories of intermediates: ‘0 to +2,’ ‘+3,’ and ‘Extended.’ (**Fig. 3B**, right brackets). The signal of each intermediate category was then quantified and fit to observe these intermediate signals over time.

**Fig. 3.**
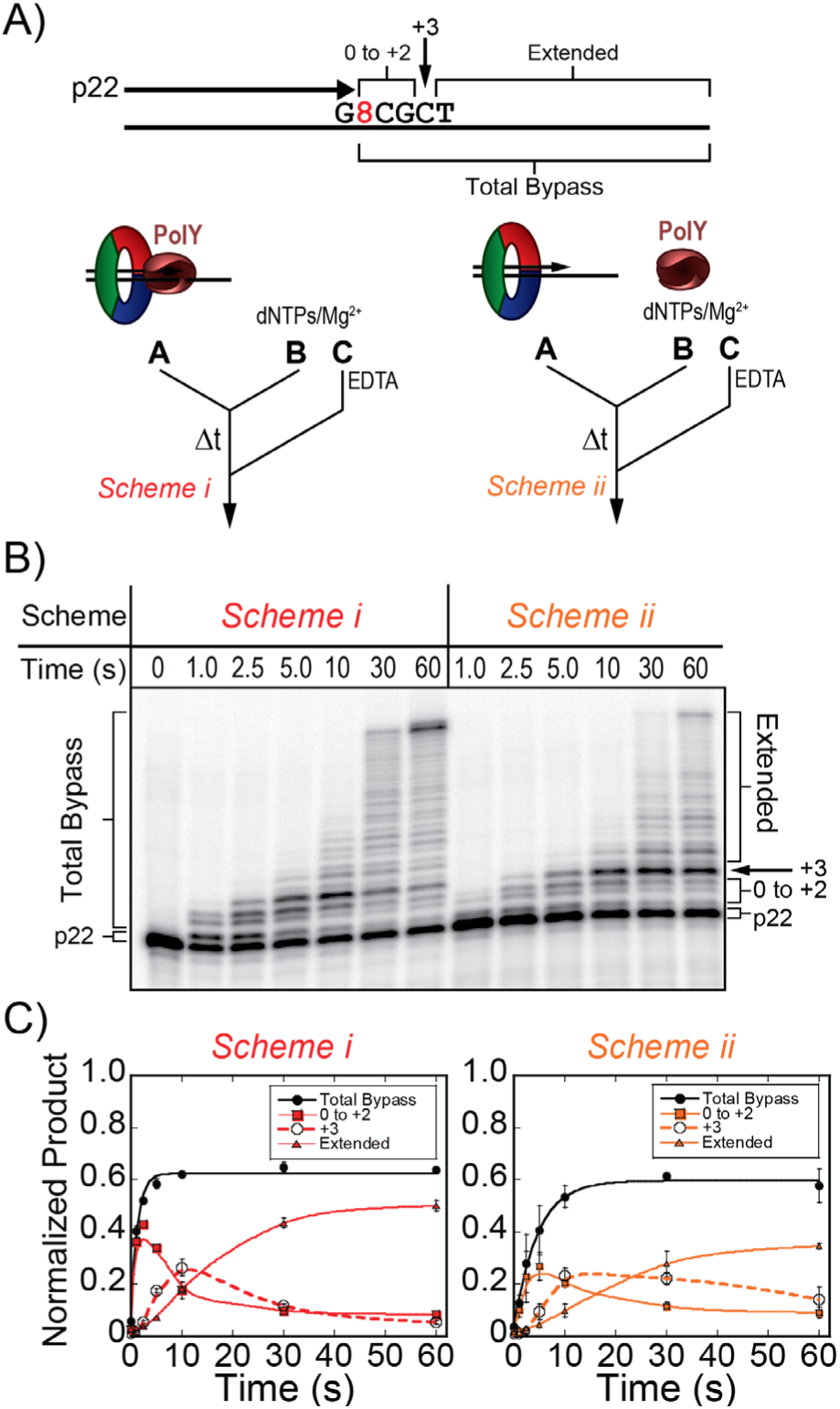
Kinetics of lesion bypass by the YHE complexes demonstrate accumulation of the +3 intermediate. A) Pre-steady-state lesion bypass assays were performed on substrates containing the stalled primer, p22 (−1), by *Schemes i* (red) and *ii* (orange). B) Products were resolved by denaturing PAGE and C) quantified for ‘Total Bypass’ (black -●-, solid line fit to **Equation 1**), or ‘0 to +2’ (−■-, smoothed solid line), ‘+3’ ‘(−∘-, smoothed dashed line), and ‘Extended’ (−▲-, smoothed solid line) intermediates. Error bars represent standard deviation from three independent replicates of each time point.

In reactions where PolY is pre-loaded with the DNA substrate and accessory factors (**Fig. 3C**, *Scheme i*), total bypass proceeds with a *k*_*obs*_ of 0.81 ± 0.11 s^-1^. This rate is representative of the catalytic rate of pre-loaded PolY across 8-oxoG. When PolY must first be recruited to the stalled DNA substrate upon initiation (**Fig. 3C**, *Scheme ii*), the apparent rate of bypass is slowed 4-fold, with a *k*_*obs*_ of 0.22 ± 0.02 s^-1^. This observed rate is likely limited by the rate of association of PolY to DNA:PCNA123 prior to the catalytic step. In these YHE schemes, the kinetic trace of the +3 intermediate accumulates (**Fig. 3C**, dashed traces) and reaches a maximal normalized product of over 0.20 (at 10 seconds), suggesting that extension by the YHE complexes is impaired at the +3 position. When lesion bypass reactions are performed by *Scheme i* using abasic and undamaged DNA templates, this intermediate is not apparent (**Supplemental Fig. S3A**). Also, on an alternative 8-oxoG template containing a different sequence context beyond the lesion (8TAC vs 8CGC), the YHE still yields an enriched +3 intermediate after 10 seconds (**Supplemental Fig. S3B**, arrow). Taken together, these results indicate that the +3 intermediate is characteristic of PolY bypass of 8-oxoG and is not an artifact of sequence context.

Considering the disappearance of the +3 intermediate and increased extension observed in the steady-state SHE reactions (**Fig. 2B and C**), it follows that addition of PolB1 into the kinetic lesion bypass assay would reduce accumulation of the +3 intermediate. To examine this, PolB1 was introduced to the pre-steady-state assay schemes through syringe B or A, opposite the location of PolY (**Fig. 4A**, *Scheme iii*, or **Fig. 4B**, *Scheme iv*), then initiated and quenched as before. When PolY is pre-loaded in syringe A, and PolB1 is introduced upon initiation in syringe B (**Fig. 4A**), lesion bypass proceeds at a similar rate (*k*_*obs*_ = 0.72 ± 0.08 s^-1^) compared to when PolY is pre-loaded alone (**Fig. 3C**, *Scheme i*). *Schemes i* & *iii* are both initiated with a pre-loaded PolY, thus quantification of ‘Total Bypass’ by these schemes is representative of the catalytic step of lesion bypass by PolY. However, in response to DNA lesions, the TLS Pol must access the primer terminus after stalling of the HiFi Pol. To investigate this step, pre-steady-state assays were performed by *Scheme iv* which simulates a stalled B1HE that has encountered an 8-oxoG lesion. Wild-type PolB1 is completely stalled by the 8-oxoG lesion (**Fig. 1C**), so ‘Total Bypass’ products by this scheme require catalysis by PolY. Therefore, in *Scheme iv*, the observed rate of synthesis would be limited by rate of the first hand-off from PolB1 to PolY. Indeed, the observed rate of lesion bypass by *Scheme iv* (*k*_*obs*_ = 0.077 ± 0.009 s^-1^) is about 10-fold slower than *Schemes i & iii* which contain a pre-loaded PolY. This rate is also 3-fold slower than the rate of lesion bypass when PolY is recruited to an unoccupied clamp (**Fig. 4B**, *Scheme iv* vs *ii*), indicating that the presence of a pre-loaded PolB1 slows PolY access to the stalled primer terminus. Observed rates of lesion bypass for *Schemes i - iv* are compared in **Figure 4C**.

**Fig. 4.**
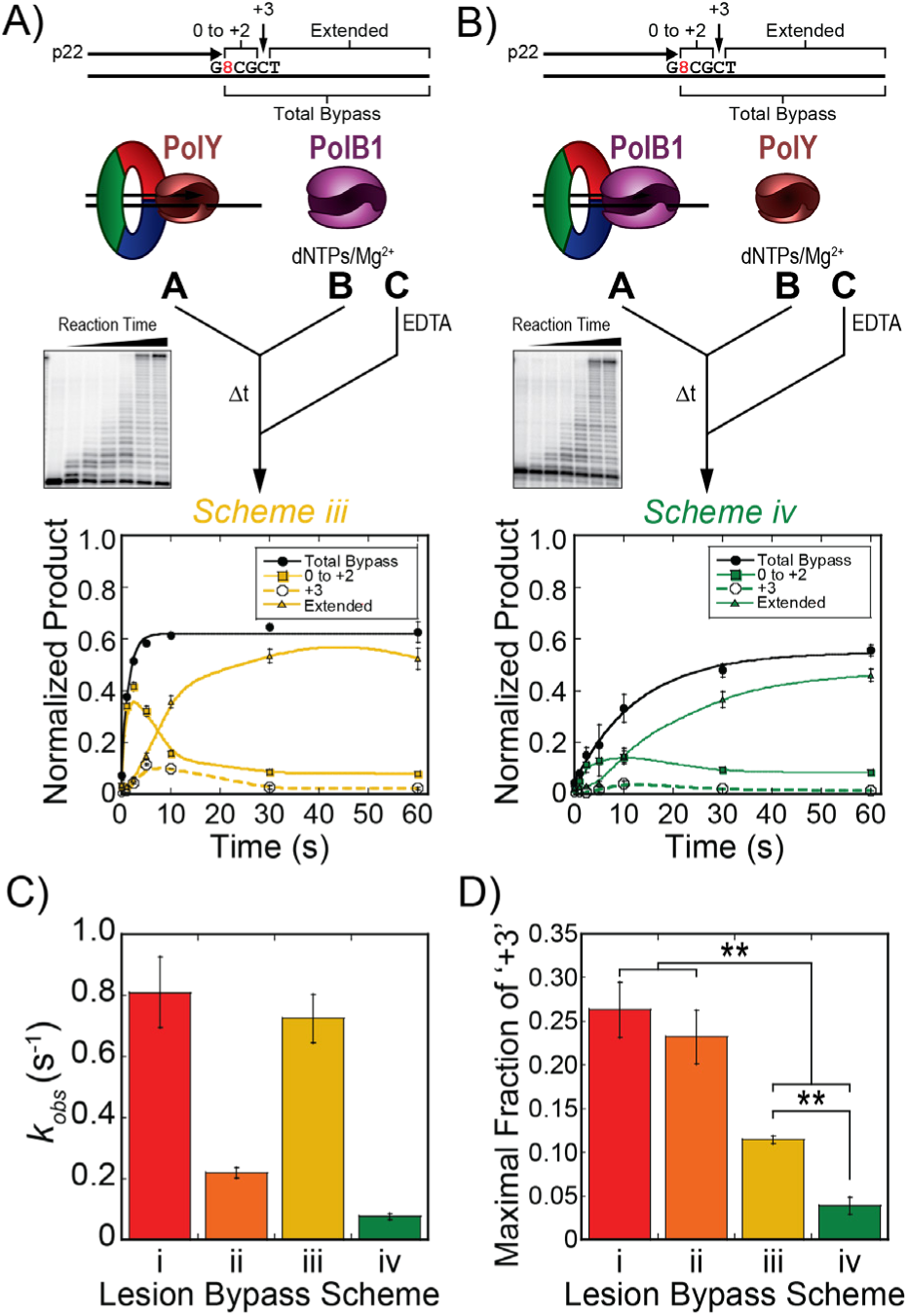
Kinetics of lesion bypass by the SHE complexes yield less +3 intermediate. Pre-steady-state lesion bypass assays were performed by A) Scheme iii (yellow) and B) Scheme iv (green). Products by Schemes iii & iv were quantified for ‘Total Bypass’ (black -●-, solid line fit to **Equation 1**) and intermediate products ‘0 to +2’ (−■-, smoothed solid trace), ‘+3’ (−∘-, smoothed dashed trace) and ‘Extended’ (−▲-, smoothed solid trace) were traced. Error bars represent standard deviation from three independent replicates of each time point. C) The observed rate constants of lesion bypass for Schemes i - iv were plotted (colors correspond to the respective schemes). Error bars indicate error of the regression fit. D) The maximal fraction of the ‘+3’ intermediate products from lesion bypass of Schemes i - iv (dashed traces) were plotted (colors correspond to the respective schemes). Error bars represent standard deviation from three replicates for each scheme.

Recently, two novel accessory subunit proteins, PBP1 and PBP2, were characterized and shown to influence the activity of PolB1 (41). PBP1 contains an acidic tail which limits strand displacement activity by PolB1 during synthesis of Okazaki fragments, whereas PBP2 moderately improves catalytic activity. Given their effects on PolB1, these PBPs were included in pre-steady-state kinetic assays on undamaged DNA templates to determine whether they have an effect in regulating or displacing PolB1 upon encountering a lesion. When both PBP1 and PBP2 (400 nM) are pre-loaded with PolY in *Scheme i*, there is no apparent change in the rate of PolY catalysis (**Supplemental Fig. S4A**). This suggests that the PBPs have no observable effect on PolY activity. Similar kinetic assays were performed using PolB1 in place of PolY (later referred to as *Scheme v* in **Fig. 6B**). By this scheme, reactions containing PBP1 demonstrate an inhibitory effect on the rate of catalysis by PolB1, while PBP2 alone improves PolB1 activity, as previously noted (**Supplemental Fig. S4B**).

**Fig. 5.**
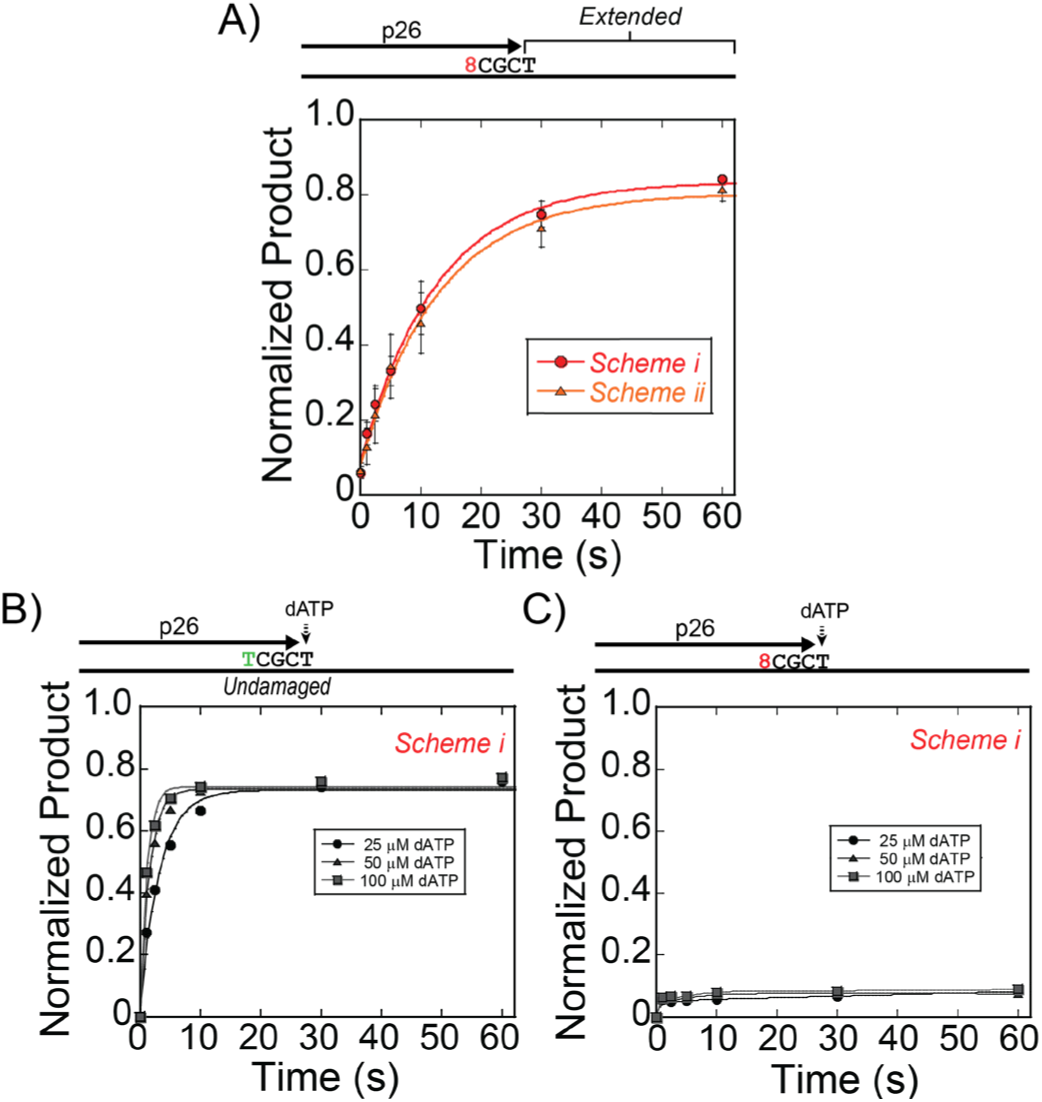
PolY conducts slow extension from the +3 intermediate. A) Pre-steady-state extension assays were performed on DNA substrates containing primer p26 (+3) by Schemes i & ii. Quantification of ‘Extended’ products were fit to Equation 1 to obtain the observed rate constants for Scheme i (red, -●-) and ii (orange, -▲-). B) Single nucleotide extension assays were performed with increasing concentrations of dATP by Scheme i on an undamaged or C) and 8-oxoG damaged template from the p26 (+3) primer as indicated in the legends. Quantification of ‘Extended’ products were fit to Equation 1 to obtain the observed rate constants at indicated concentrations of dATP.

**Fig. 6.**
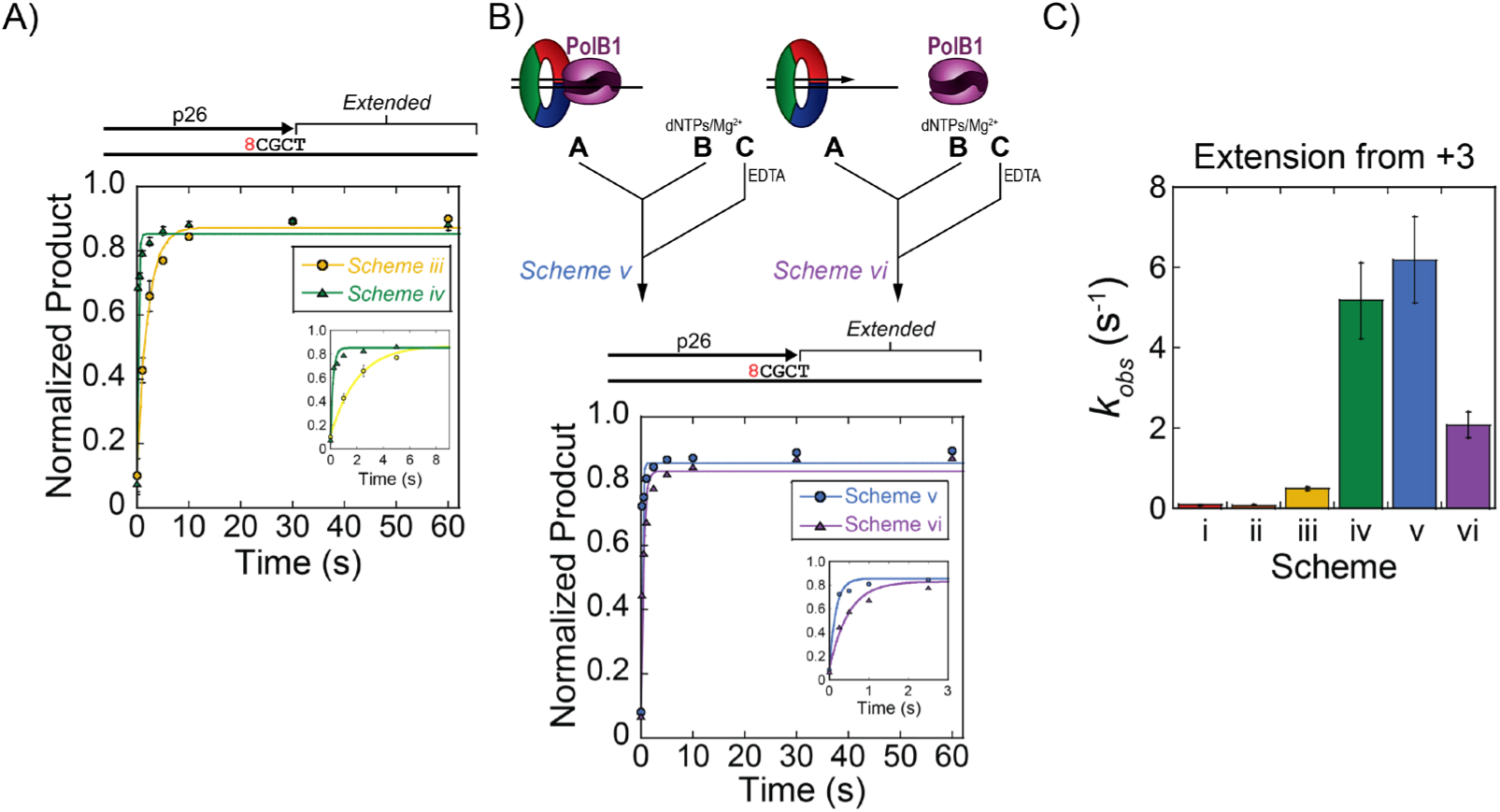
PolB1 conducts rapid extension from the +3 intermediate position. Pre-steady-state extension kinetics were performed on a DNA substrate containing primer p26 (+3) by Schemes iii (yellow, -●-) and iv (green, -▲-) containing PolB1 and PolY, or B) Schemes v (blue, -●-) and vi (purple, -▲-) containing PolB1 alone. Error bars represent standard deviation from three independent replicates of each time point. Quantification of ‘Extended’ products were fit to Equation 1 to obtain the observed rate constant for extension. Insets show the trace of ‘Extended’ product at a shorter time scale. C) Observed rate constants for extension from +3 were plotted; error bars indicate error of the regression fit.

From its apparent inhibition of PolB1 activity, we hypothesized that PBP1 may destabilize PolB1 affinity to DNA. Upon PolB1 encountering a template lesion, PBP1-mediated displacement of PolB1 would allow faster access of PolY to the stalled primer. To investigate this, lesion bypass kinetic assays were performed by *Scheme iv* in the presence of PBP1 pre-loaded with PolB1 in syringe A. However, this did not yield any apparent change in the observed rate of exchange from PolB1 to PolY for lesion bypass (**Supplemental Fig. S4 C and D**). The similarity in these rates suggests that PBP1 does not regulate PolB1 activity by promoting its displacement from DNA, and these accessory subunits are not directly involved in the Pol hand-off during ‘on-the-fly’ TLS.

To compare the accumulation of the +3 intermediate in all lesion bypass kinetics schemes (**Figs. 3C, 4A, and 4B**, dashed traces), we plotted the maximal signal of the +3 intermediate in the kinetic traces from each Pol complex (**Fig. 4D**). In schemes containing PolB1 (*iii* & *iv*), the maximal fraction of the +3 intermediate (0.11 ± 0.004 and 0.04 ± 0.01, respectively) is significantly less than the intermediate signal from reactions containing PolY alone (*Schemes i* & *ii*; 0.26 ± 0.03 and 0.23 ± 0.03, respectively). Despite the slower overall rate of lesion bypass in *Scheme iv*, the reduced fraction of +3 intermediate indicates that extension from the intermediate position is still fast.

Again, this shows that PolY is capable of rapid lesion bypass up to the +3 position, but additional extension is impaired and results in accumulation of the +3 intermediate which is relieved by the presence of PolB1.

### Slow extension by the YHE complexes from the +3 intermediate indicates a perturbation in PolY catalytic activity at this position

The peculiar features of this +3 lesion bypass intermediate prompted us to investigate the kinetics of extension from this position by the different Pol complexes. Reactions were assembled in the RQF-3 as described above, except a DNA substrate containing primer p26 (+3), which corresponds with the observed intermediate, was utilized. All products beyond the primer were quantified as ‘Extended’ products (**Fig. 5A**). For reactions containing PolY alone (*Schemes i* & *ii*), the apparent rates of extension are equivalent for *Schemes i* & *ii*, regardless of PolY syringe position (*k*_*obs*_ = 0.080 ± 0.008 s^-1^ and 0.078 ± 0.008 s^-1^, respectively). The similar rates from these pre-loading schemes suggest that extension by PolY from the +3 position is not limited by its rate of association to the DNA substrate. This contrasts with the observed rates obtained for lesion bypass for *Schemes i* & *ii* (**Fig. 3c**). These YHE extension rates are about 2.8-fold slower than the apparent rate of association for PolY (represented by lesion bypass in *Scheme ii*), indicating that the rate of PolY catalysis is limiting when extending from the +3 intermediate position after 8-oxoG. We considered the reduced rate of extension to be limited by a decrease in DNA binding affinity of PolY to the +3 extension substrate. This circumstance would effectively reduce the formation of active YHE complexes and yield a slower apparent rate of extension. However, as determined by EMSA, there was no apparent change in PolY binding affinity for the +3 intermediate substrate relative to an to an 8-oxoG damaged substrate with a primer at the stalled position, nor to a substrate containing an abasic site lesion or undamaged base (**Supplemental Fig. S5**). This suggests that slow extension of the YHE from the +3 intermediate is not a result of destabilized DNA binding, rather it is limited by a slow elementary step which follows PolY association with the DNA substrate.

To investigate the kinetic parameter of PolY that is affected at this position, we measured the observed rates of single nucleotide incorporation by S*cheme i* as a function of dATP concentration, the next nucleotide for extension from the +3 position. On an undamaged DNA template (t52_Und_), *k*_*obs*_ increases with elevated [dATP], as expected (**Fig. 5B**). However, dATP incorporation from this position on the 8-oxoG damaged template results in a negligible amount of extension product and did not allow for calculation of a *k*_*obs*_ within the measured time scale (**Fig. 5C**). Though we cannot quantitatively compare observed rates of extension, these data qualitatively and obviously indicate a major disruption of an elementary polymerization step by PolY at this +3 position and provide a kinetic basis for accumulation of the intermediate downstream of the 8-oxoG lesion.

### PolB1 performs rapid extension from the +3 intermediate position

As observed in **Figure 1D**, PolB1 extension activity is highest when starting from the +3 position (purple circles), indicating kinetic efficiency for extension by PolB1. In order to investigate the role of PolB1 in extension from this intermediate position, we evaluated the kinetics of extension from the +3 position by *Schemes iii & iv*. Extension by *Scheme iii* simulates a condition that examines a switch from PolY to PolB1 at the +3 intermediate. This observed rate of extension (0.49 ± 0.05 s^-1^, **Fig. 6A**, *Scheme iii*) is 6.1-fold faster than extension by the YHE complex (**Fig. 5**, *Schemes i* & *ii*). Since PolY is greatly inhibited at this position (**Fig. 5C**), the faster extension rate in *Scheme iii* is attributed to the activity of PolB1 and is representative of the second hand-off from TLS PolY to HiFi PolB1 for extension. In *Scheme iv*, PolB1 is already pre-loaded on the +3 extension substrate and yields a fast *k*_*obs*_ of 5.2 ± 0.9 s^-1^ (**Fig. 6A**, *Scheme iv*). This represents the catalytic rate of extension by a pre-loaded PolB1 and is 65-fold faster than extension by the YHE complexes from the +3 position (**Fig. 5A**, *Schemes i* & *ii*). Together, these data indicate that PolB1 can access the 3’-end from PolY at the +3 position faster than PolY can perform additional extension, and PolB1 is capable of rapid and efficient extension from this position.

Extension kinetics reactions were also performed with PolB1 alone introduced from either syringe A or B (**Fig. 6B**, *Schemes v* & *vi*, respectively). Extension by *Scheme v* resembles *Scheme iv*, containing a pre-loaded PolB1, and yields a similar rate of extension (*k*_*obs*_ = 6.2 ± 1.2 s^-1^). When PolB1 must be recruited to the substrate upon initiation (*Scheme vi*), the rate of extension is also relatively fast (*k*_*obs*_ = 2.1 ± 0.3 s^-1^). This observed rate likely represents the rate of association for PolB1 to an unoccupied PCNA123:DNA complex. These assays highlight the rapid extension activity of PolB1 from the +3 position. These rates (*Schemes v* & *vi*) are orders of magnitude faster than extension by PolY (*Schemes i* & *ii*) and provide kinetic evidence for the position where PolB1 resumes HiFi extension activity after PolY bypass of 8-oxoG. Observed rates of extension for *Schemes i - vi* are compared in **Figure 6C**.

## Discussion

In this study, we have identified a novel +3 intermediate for PolY after bypass of an 8-oxoG lesion, which results from a perturbation in polymerase activity at that position. In the presence of PolB1, accumulation of the +3 intermediate is reduced and rapidly extended into longer products. Altogether, this points to an ‘on-the-fly’ TLS mechanism where the first hand-off (from stalled PolB1 to PolY) occurs at the -1 position (**Fig. 7A to B**). PolY then performs TLS and extends to a position 3 base pairs beyond the lesion, where it becomes catalytically inefficient for extension to the +4 position (**Fig. 7C**). From this +3 position, the catalytic inefficiency PolY allows for the second hand-off to restore PolB1 activity (**Fig. 7D to E**). This marks a distinct position beyond an 8-oxoG lesion for the second Pol hand-off which resumes HiFi replication and effectively limits error-prone TLS activity.

**Fig. 7.**
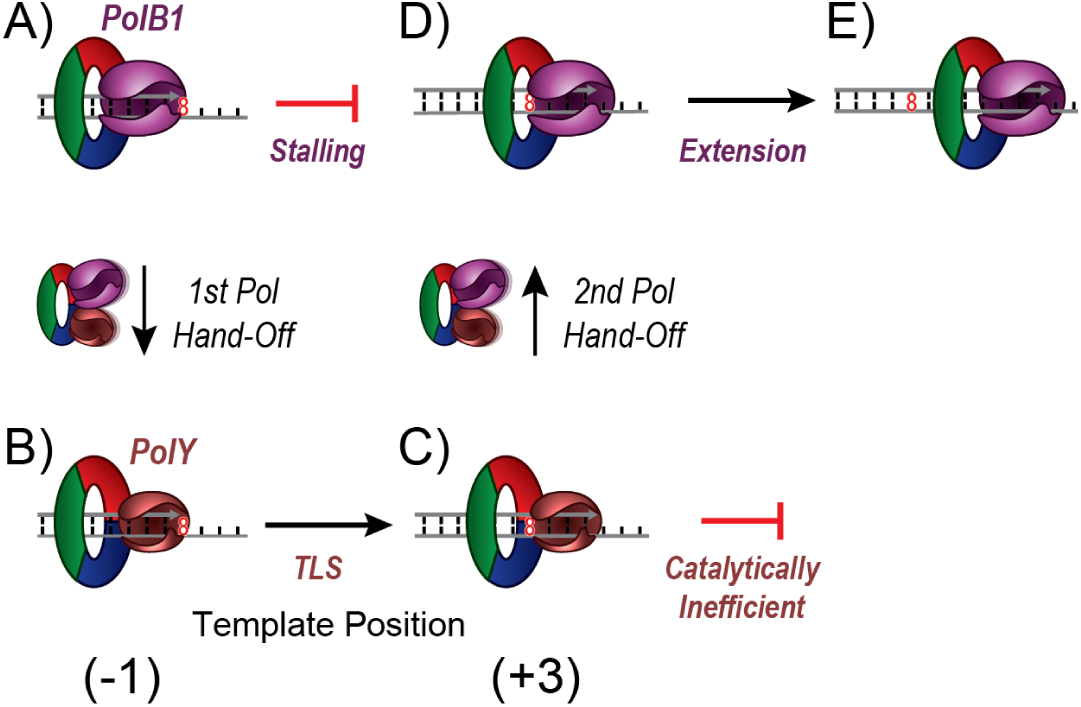
Proposed mechanism for DNA hand-offs between PolB1 and PolY in response to an 8-oxoG lesion. A) Upon encountering 8-oxoG on the template strand, PolB1 stalls. B) Individual contacts with PCNA123 mediate the first Pol hand-off from PolB1 to PolY. C) PolY then performs translesion synthesis to the template position three base pairs beyond the 8-oxoG lesion, where it becomes catalytically inefficient for additional synthesis. D) The second Pol hand-off from PolY to PolB1 is then mediated through PCNA123 to re-establish HiFi PolB1 for E) extension and effective bypass of the lesion.

Previous kinetic and structural studies of PolY bypass and extension across 8-oxoG did not identify the +3 intermediate (24,25,28). Accumulation of this intermediate was first apparent to us upon inclusion of spDNA trap in steady-state lesion bypass assays, which effectively amplifies sites of catalytic inefficiency during replication. In prior studies without trap DNA, multiple enzyme turnovers and distributive extension from the +3 position may have proceeded too quickly for its accumulation to be apparent. These studies also primarily focus on the kinetic steps of lesion bypass, translocation, and extension up to the +1 position by PolY; its activity was not evaluated further downstream of 8-oxoG. Finally, prior evaluations were also performed without a processivity clamp, which is a critical accessory factor that optimizes and mediates Pol and other enzymatic activities (39,42,43). Our study suggests that PCNA123 mediates the complementing activities of PolY and PolB1 not only upon encountering the lesion, but also after the lesion has been bypassed. In response to other DNA lesions, the kinetics of lesion bypass by PolY have already been characterized (44-49). In some of the previous analyses, ‘pause sites’ were noted during PolY lesion bypass, indicating points of inefficient catalysis and extension from these lesions. However, in the absence of RFC, PCNA123, and PolB1, these pause sites cannot be directly correlated with the position of the second hand-off after lesion bypass. This prompts additional investigation into how PolY and other TLS Pols are limited in extension after bypass of various DNA lesions, especially in the presence of a processivity clamp and HiFi Pols.

Interestingly, *E. coli* Pol IV (or DinB, a homolog of PolY) (22) demonstrates a similar +3 intermediate following bypass of N^2^-dG-peptide adducts (50). Also, a Pol IV variant led to identification of a +3 intermediate in response to an N^2^-furfuryl-dG lesion, with a drastic reduction in catalytic efficiency from the intermediate position (51). Both studies suggest the potential for structural impediments, or communication, between a bypassed template lesion and the TLS Pol active site once it has translocated to a certain position downstream, but the basis of this disruption has not been explored. We hypothesize that the observed catalytic inefficiency at the +3 intermediate corresponds with a disruption in translocation of PolY from this position. A series of positively charged residues of the PolY LF domain interact with the DNA backbone and are implicated in facilitating PolY translocation (25). During extension from a bypassed lesion, an altered base with modifications that are oriented toward the LF may sterically disrupt interactions between template DNA and the residues involved in translocation. A disruption in translocation would prevent the Pol active site from accessing the next template base and interrupt binding of an incoming nucleotide, ultimately weakening the apparent efficiency of successive catalytic extensions. The apparent conservation of the +3 intermediate for TLS Pols from both *E. coli* and *Sso* strengthens the hypothesis that the LF domain of PolY (and other DinB homologs) can sense bypassed lesions after translocating to a certain position downstream. Activation of this ‘pinky trigger,’ along with complementation of extension activity by a HiFi Pol and mediation through a processivity clamp, serves to minimize the extent of error-prone DNA synthesis by TLS Pols and restore HiFi Pol activity.

Previously, under conditions similar to the current study, we measured the rate of association of PolY to a pre-loaded B1HE using pre-steady-state FRET (13). This apparent rate (0.8 ± 0.1 s^-1^) is over 10-fold faster than the apparent rate of lesion bypass by the same pre-loading approach evaluated in *Scheme iv* (0.077 ± 0.009 s^-1^). In that same study, we also observed that PolB1 was not displaced upon PolY association within the measured timescale, indicating that both Pols were simultaneously bound to the clamp and formed a concerted PCNA tool belt (13). Combined with our current findings, this suggests that PolY may rapidly associate with the stalled B1HE to form a SHE complex, but the rate of the first Pol switch within the complex (involving transient dissociation of PolB1 and PolY binding to the stalled substrate) is slower and rate limiting for lesion bypass by PolY.

Interestingly, the apparent rate for the first Pol hand-off to PolY for TLS (0.077 ± 0.009 s^-1^) is 6.4-fold slower than the second switch to PolB1 for extension (0.49 ± 0.05 s^-1^; **Fig. 4a** *Scheme iv* vs **Fig. 6a** *Scheme iii*). This suggests that PolB1 is more stably bound to DNA relative to PolY, which is consistent with previous Pol:DNA binding affinity studies (52), yet these binding affinities have not been evaluated in the presence of PCNA123. The apparent difference in hand-off rates observed in this study indicates that the fundamental basis for each hand-off is molecularly unique, and additional investigation is also required to obtain direct insight into the kinetic and structural mechanisms of these Pol hand-off events.

After an ‘on-the-fly’ TLS event and resumption of HiFi Pol activity, the TLS Pol may remain bound to the clamp during replicative extension. This configuration allows TLS to be transiently available for subsequent encounters with DNA lesions. Yet, in the context of a more complex multi-equilibria system, dynamic exchange of HiFi Pols and other DNA processing enzymes has been observed (53-55). PCNA123 has also been shown to coordinate an alternate arrangement of enzymes, optimizing their combined activities for Okazaki fragment maturation (56). This indicates that an effective cellular response would require dynamic configurations of various cellular tools in order to faithfully maintain the genome. Therefore, the mechanisms of Pol hand-offs may include aspects of distributive Pol exchange. However, this does not eliminate the possibility of switching through direct contacts between Pols by way of a transiently concerted PCNA tool belt complex, especially considering the network of direct interactions among TLS Pols (57,58) and the sharing of accessory subunits between eukaryotic TLS Polz and HiFi Pold (59-61).

Characterizing the intricacies of ‘on-the-fly’ TLS and its hand-off mechanisms is integral to our understanding of this early DNA damage response. Though this damage tolerance pathway benefits the cell by relieving stalled replication, unregulated TLS activity can detrimentally contribute to mutagenesis and resistance to DNA-damaging cancer therapies. Recent advances have addressed these drawbacks through identification of small molecule inhibitors which target Pol-Pol interactions and/or disrupt mutagenic TLS activity, supplementing the effectiveness of current chemotherapeutic drugs (62-68). These studies provide a promising new approach for mitigating the role of TLS in chemoresistance and provide an exciting incentive for furthering our understanding of the kinetic and structural foundations of TLS.

## Materials and Methods

### Protein expression and purification

Wild type Sso PolB1 (38), PolY (13), and RFC (39), were expressed and purified as previously described. The wild-type variant of PolB1 was used in these studies, as exonuclease deficient PolB1 has been shown to bypass 8-oxoG lesions (28). His-tagged subunits of heterotrimeric PCNA123 were expressed and purified individually and reconstituted into the heterotrimer complex as previously described (39).

The DNA sequence for PolB1 binding protein 1 (PBP1; SSO0150) and PBP2 (SSO6202) were amplified from Sso genomic DNA (primer sequences in Supplemental Table 1). The amplified genes were cloned into pET30a using XhoI and NdeI restriction sites. The proteins were expressed in Rosetta2 DE3 E. coli cells by autoinduction (40), followed by lysis and purification of the PBPs as previously described (41).

### Steady-state lesion bypass and extension assays

DNA primers (Sigma) were radiolabeled at the 5’-end with 32P using the manufacturer recommended protocol for T4 PNK (New England Biolabs) and ATP [g-32P] (Perkin Elmer). Radiolabeled primers were individually mixed with the DNA template containing an 8-oxoG lesion at the 23rd base from the 3’ end (t528; Fig. 1A and Supplemental Table 1). Substrates were annealed at a 1.2:1 primer:template ratio in a thermocycler (BioRad) by incubation for 10 minutes at 95 oC, followed by a decrease to 20 oC at a rate of 1 oC/min. Annealed substrates were pre-loaded with different Pol HE complexes in reaction buffer (20 mM Tris OAc (pH 7.5), 100 mM KOAc) with PCNA123, RFC, PolB1 and/or PolY. ATP was added, followed by incubation at 60 oC for 5 min to promote loading of PCNA123 to the DNA substrate. Reactions were initiated upon addition of dNTPs, Mg(OAc)2, and salmon sperm DNA (spDNA; Invitrogen) as an enzyme trap, followed by quenching with 100 mM EDTA after 3 minutes. The final reaction concentration of each component is indicated in Figure 1A.

Reaction products were resolved by denaturing polyacrylamide gel electrophoresis (PAGE) through either a 10 cm gel (20% polyacrylamide/6M urea/25% formamide/1x TBE) or a 38 cm DNA sequencing gel (15% polyacrylamide/6M urea/25% formamide/1x TBE). Resolved products were exposed to a phosphor screen for a minimum of 4 hours and scanned with a Storm 820 phosphorimager (GE Healthcare Life Sciences). The product signals were quantified using ImageQuant software (v5.0, GE Healthcare Life Sciences).

### Pre-steady-state lesion bypass and extension kinetics assays

For pre-steady-state lesion bypass kinetics assays, reactions were performed using a DNA substrate containing the 8-oxoG damaged template (t528) annealed to primer p22 (−1). This ‘stalled’ DNA substrate was pre-mixed with reaction buffer, PCNA123, and RFC. ATP was added followed by incubation for 2 minutes at 60 oC and loading into sample loop A of an RQF-3 rapid-quench instrument (KinTek). The RQF-3 was connected to a circulating water bath at 40 oC. This temperature was chosen based on a balance of the temperature limitations recommended by the instrument manufacturer and the optimal reaction temperature for the enzymes (27). Reactions were initiated upon addition of dNTPs and Mg(OAc)2 from sample loop B, followed by quenching with EDTA from syringe C after indicated time points. PolY and PolB1 were introduced into lesion bypass kinetics assays from different syringe positions as outlined by pre-loading Schemes i - iv. Final concentrations of all reaction components are the same as outlined in Fig. 1A. Products were resolved by denaturing PAGE, exposed, and scanned as described above. In order to obtain the observed rate constant for lesion bypass, the normalized signals for total bypass products were fit to according to Equation 1:

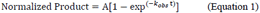

where A is the reaction amplitude, kobs is the observed rate constant, and t is reaction time (Kaleidagraph, v4.5). Signals of other reaction intermediates (defined by brackets) were also plotted and fit to smooth traces for visual comparison of intermediate signal magnitudes.

For pre-steady-state extension kinetics assays, reactions were performed using a DNA substrate containing an 8-oxoG damaged (t528) or undamaged (t52Und) DNA template annealed to primer p26 (+3). Where indicated, single nucleotide incorporation assays were performed by initiating extension reactions with various concentrations of dATP. PolY and/or PolB1 were introduced to the extension kinetics assays from different syringe positions as outlined in pre-loading Schemes i - vi. Products were resolved by denaturing PAGE, exposed, scanned, and quantified for total extension activity as above.

### Electrophoretic Mobility Shift Assay (EMSA)

Increasing concentrations of PolY were incubated with 4 nM of the indicated radiolabeled DNA substrates in reaction buffer with 5 mM magnesium acetate and 10% glycerol. Reactions were incubated at 40 oC for 10 min and resolved through a 10 cm 4% polyacrylamide/1xTBE native PAGE gel. Gels were exposed to a phosphor screen for a minimum of 4 hours and scanned using a Storm 820 phosphorimager (GE Healthcare Life Sciences). The bound fraction of each DNA substrate was plotted as a function of PolY concentration and fit to Equation 2 to obtain the dissociation constant, Kd:

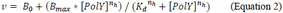

where B0 is the fraction bound at 0 nM PolY, Bmax is the fraction bound at saturation, nh is the Hill slope, and Kd is the dissociation constant for PolY binding to DNA.

## SUPPLEMENTAL DATA

Supplementary Data are available online.

## ACKNOWLEDGEMENTS

We acknowledge the Baylor Molecular Bioscience Center (MBC) for providing instrumentation and resources aiding this project. Also thanks to the Finkelstein lab for providing the bioRxiv template. https://github.com/finkelsteinlab/BioRxiv-Template

