## Supplemental for "A hand-off of DNA between archaeal polymerases allows high-fidelity replication to resume at a discrete intermediate three bases past 8-oxoguanine"

### SUPPLEMENTAL DATA

#### Supplemental Table S1: DNA substrates used in this study

***Substrate Primers***

| Name | Position relative to lesion | Sequence (5'-3') |
|---|---|---|
| p18 | -5 | GCGAGGCGAGCGCGAGCG |
| P22 | -1 | GCGAGGCGAGCGCGAGCGATAC |
| P23 | 0 | GCGAGGCGAGCGCGAGCGATACC |
| P24 | +1 | GCGAGGCGAGCGCGAGCGATACCG |
| P25 | +2 | GCGAGGCGAGCGCGAGCGATACCGC |
| P26 | +3 | GCGAGGCGAGCGCGAGCGATACCGCG |
| P41 | -1 | GCGAGGCGAGCGCGAGCGATACCGCGATCGAGTGCAAGCTT |

***Substrate Templates***

| Name | Description | Sequence (5'-3') |
|---|---|---|
| $t52_{Und}$ | Undamaged 52mer template | CGCTCCGCTCGCGCTCGCTATGTCGCTAGCTCACGTTCGAAGTACAACCTGC |
| $t52_{8}$ | 8-oxo-G template (@23) | CGCTCCGCTCGCGCTCGCTATG8CGCTAGCTCACGTTCGAAGTACAACCTGC |
| $t52_{AP}$ | Abasic template (@23) | CGCTCCGCTCGCGCTCGCTATG_CGCTAGCTCACGTTCGAAGTACAACCTGC |
| $t52_{Alt8}$ | Double 8-oxo-G template (@23 and 42) | CGCTCCGCTCGCGCTCGCTATG8CGCTAGCTCACGTTCGAA8TACAACCTGC |

8 – 8-oxoguansine; _abasic site

***Cloning Primers***

| Name | Sequence (5'-3') |
|---|---|
| SsoPBP1+NdeI FWD | ATTACATATGTCAACGAGATGGCTACCTAAG |
| SsoPBP1+XhoI REV | ATTACTCGAGTTACTCCTCTTCACTTTCTTCTTCAC |
| SsoPBP2+NdeI FWD | ATTACATATGAATACTGGATTAATATATCTTATGTCTGTTAATC |
| SsoPBP2+XhoI REV | ATTACTGCAGTCACTTCTTGTCAGTAGATTTCCTCAC |

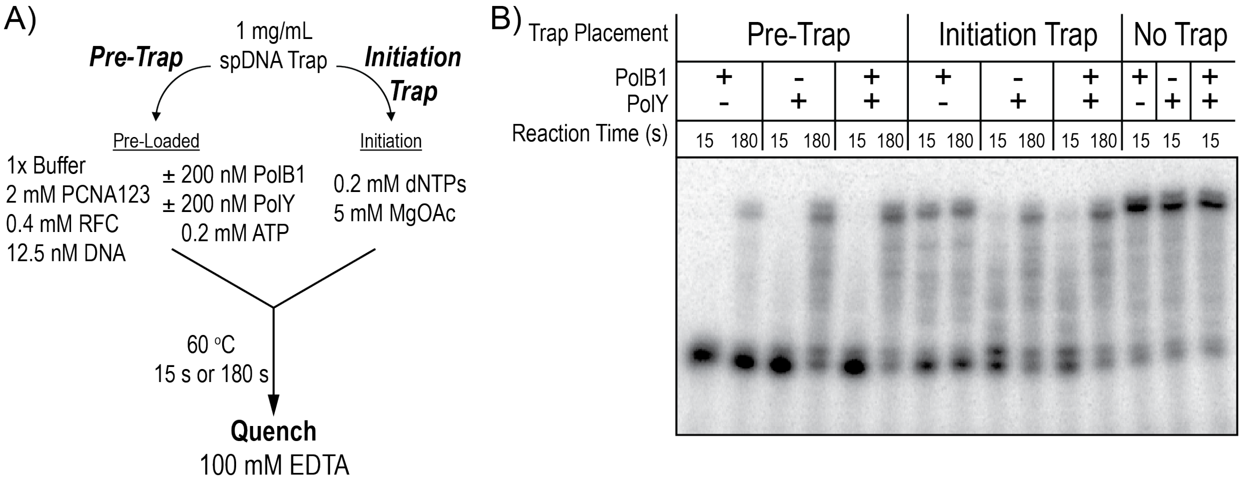


**Supplemental Figure S1: Pre-trapping with 1 mg/mL of spDNA still allows for distributive Pol activity.** A) 1 mg/mL of spDNA was introduced to the steady-state polymerase assay either by pre-mixing with the indicated Pol complex and undamaged DNA substrate ('Pre-Trap'), added upon initiation with dNTPs and $Mg^{2+}$ ('Initiation Trap'), or in the absence of trap ('No Trap'), and quenched at indicated time points. B) All 'Pre-Trap' Pol complexes demonstrate time-dependent activity, indicating that this concentration of trap still allows for distributive exchange of Pols from solution through an altered multieuilibria.

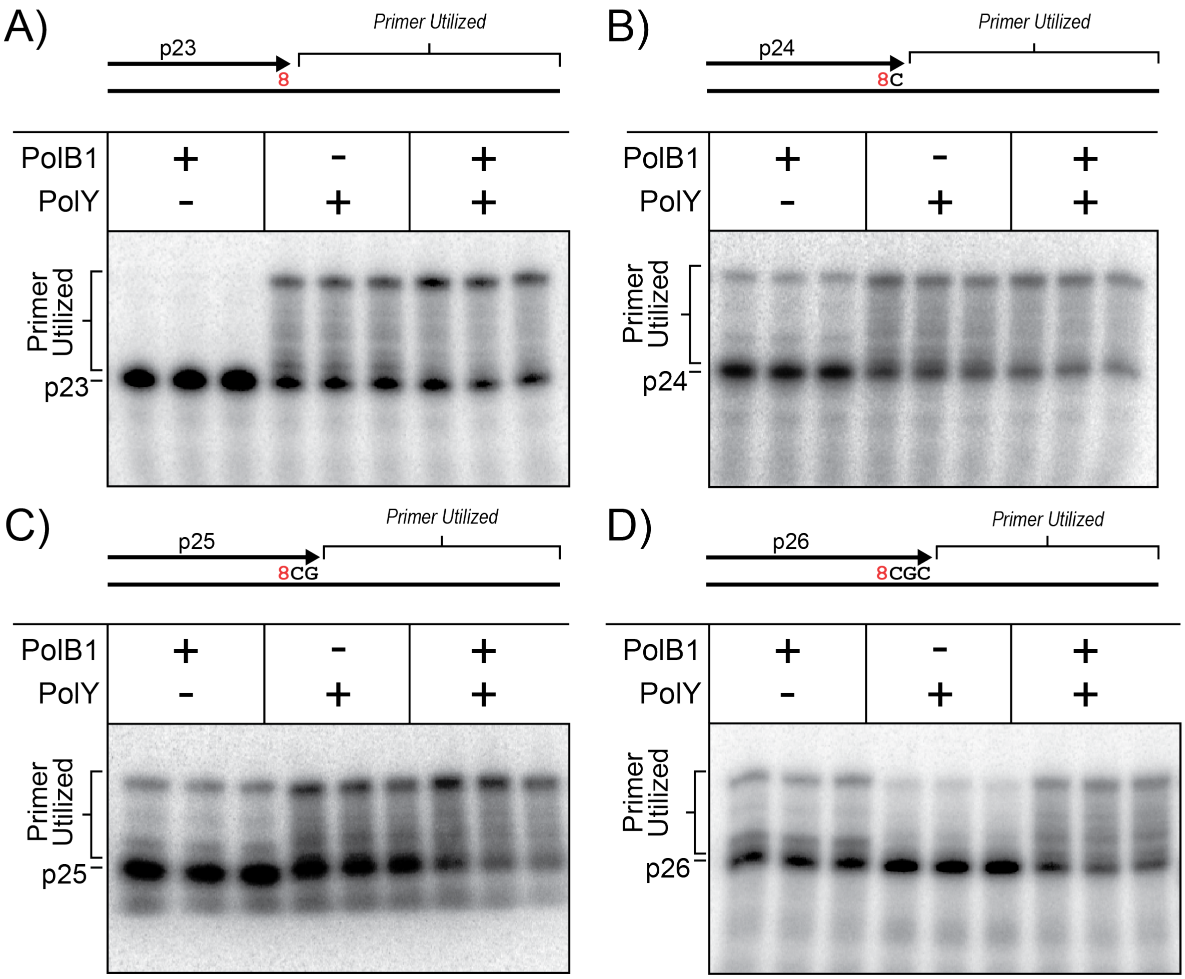


**Supplemental Figure S2: Gel images of steady-state lesion bypass by each Pol complex on progressively longer primers.** Reactions described in **Figure 1A** were performed on 8-oxoG damaged DNA substrates containing progressively longer primers A) p23, B) p24, C) p25, and D) p26. Products were quantified as a percentage of primer utilized (brackets) and plotted in **Figure 1D**.

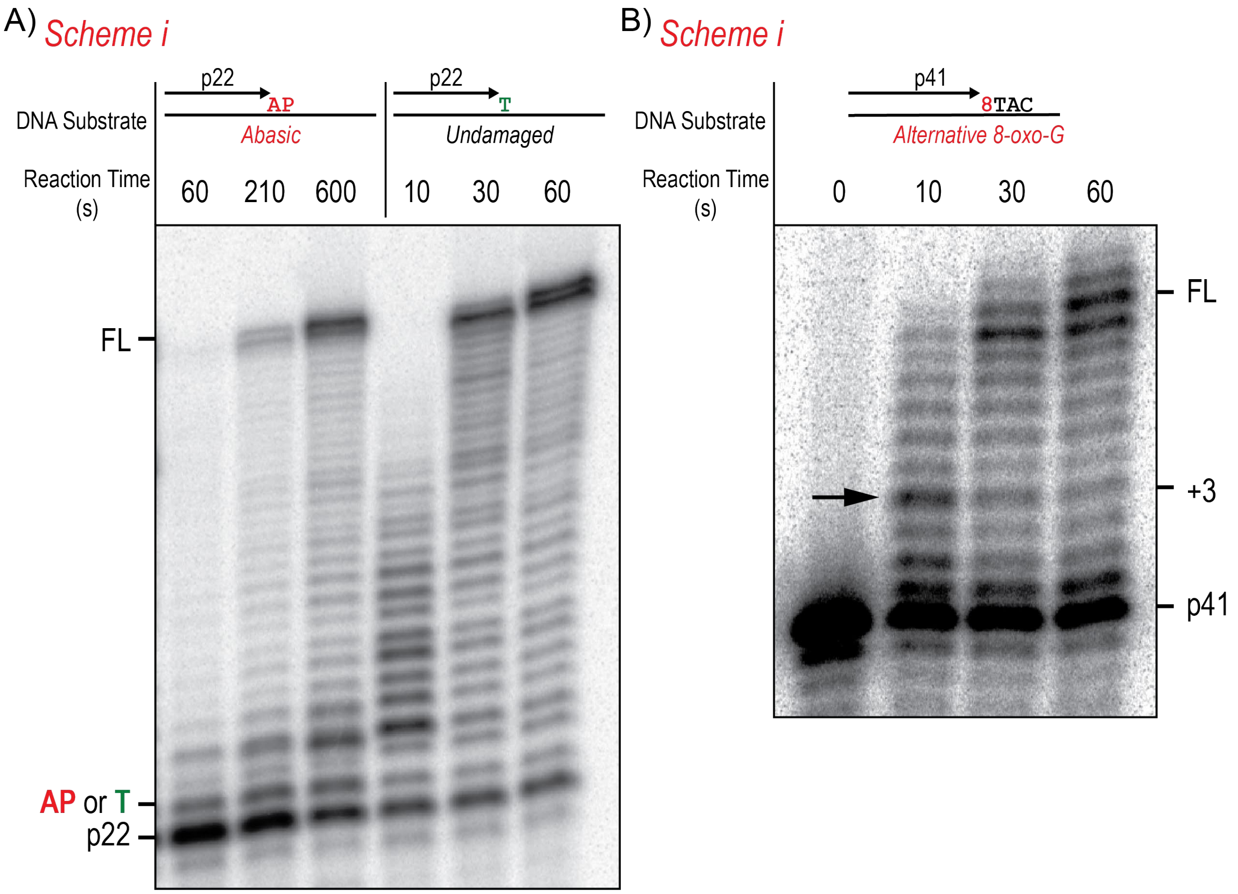


**Supplemental Figure S3: The +3 intermediate is characteristic of PolY bypass of 8-oxoG and is not an artifact of sequence context beyond the lesion.** Pre-steady-state lesion bypass kinetics reactions were performed by *Scheme i* on DNA substrates containing A) an abasic site (AP) lesion or an undamaged T. Similar reaction were also performed on an alternative 8-oxoG template with a different sequence context beyond the lesion.

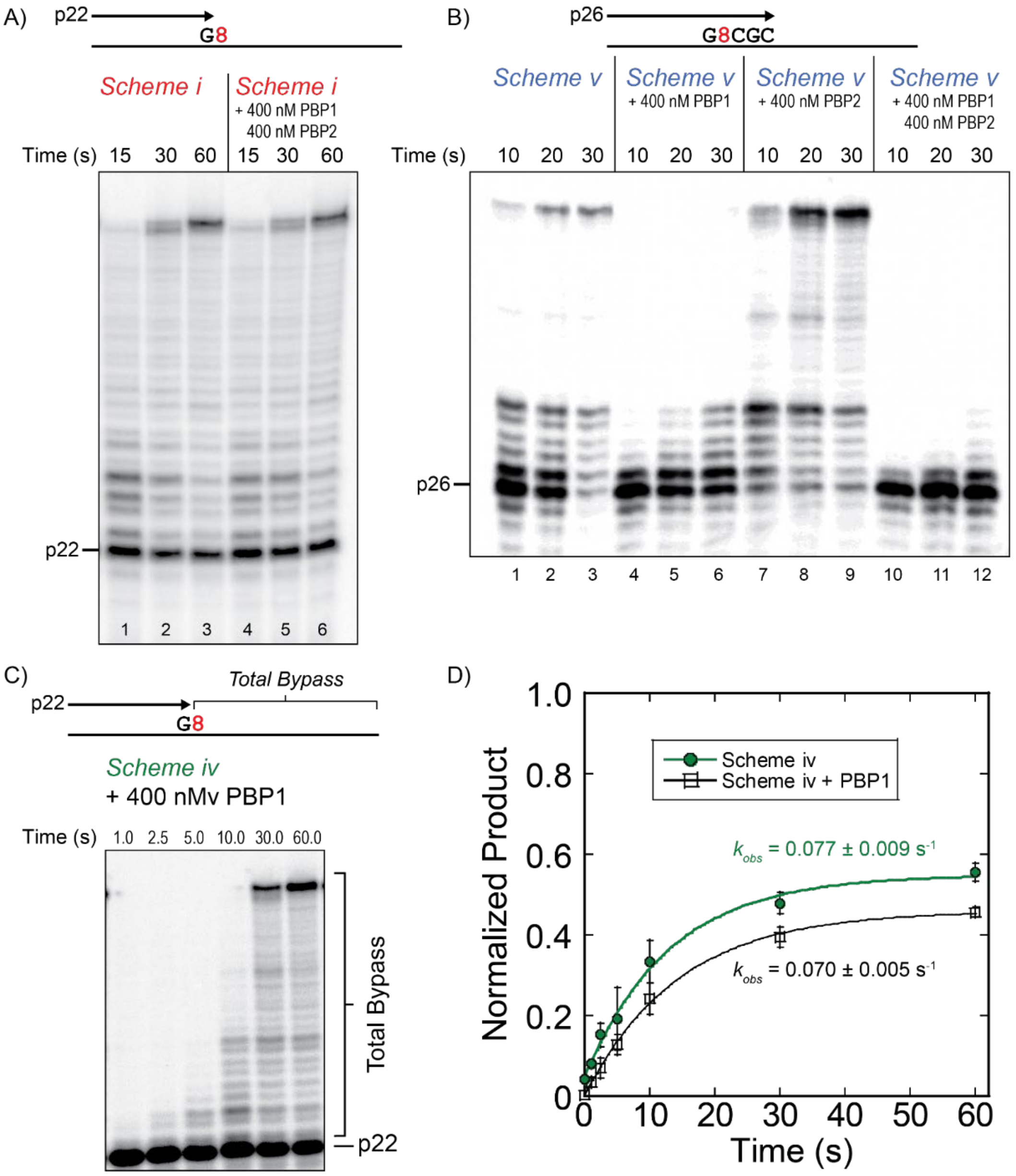


**Supplemental Figure S4: PBP1 does not increase the rate of the first hand-off from a stalled PolB1 to PolY for lesion bypass.** A) Pre-steady state lesion bypass kinetics were performed by *Scheme i* (containing pre-loaded PolY) on the indicated DNA substrate either alone (lanes 1-3) or in the presence of 400 nM PBP1 and PBP2 (lanes 4-6). B) Pre-steady-state extension kinetics assays were performed by *Scheme v* (containing pre-loaded PolB1; see **Fig. 6A**) on the indicated DNA substrate. Reactions were performed with PolB1 alone (lanes 1-3), or in the presence of 400 nM PBP1 (lanes 4-6), 400 nM PBP2 (lanes 7-9), or 400 nM of PBP1 and PBP2 (lanes 10-12). C) Similar reactions were performed by *Scheme iv* with 400 nM PBP1 pre-loaded in syringe A and resolved by denaturing PAGE. D) 'Total Bypass' products were quantified over time (black, -□-) and fit to **Equation 1** to obtain the observed rate constant and compared to the rate of *Scheme iv* in the absence of PBP1 (green, -●-).Error bars represent the standard deviation of three independent replicates for each time point.

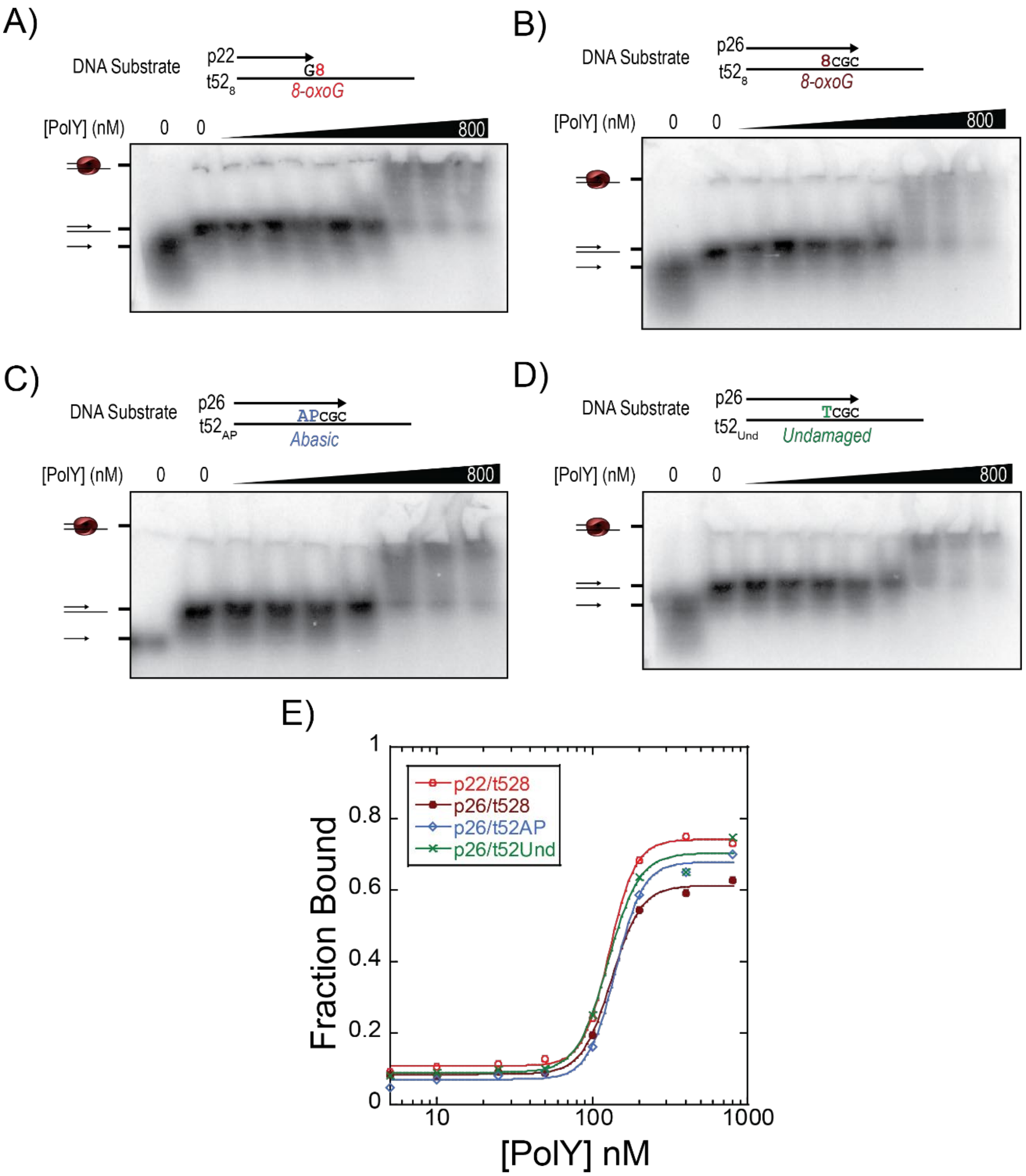


**Supplemental Figure S5: Inefficient extension of PolY from the +3 intermediate is not a result of decreased DNA binding affinity.** EMSAs were performed by incubating increasing concentrations of PolY to various DNA substrates containing A) p22/t52$_8$, B) p26/t52$_8$, C) p26/t52$_{AP}$, and D) p26/t52$_{Und}$. E) The fraction of DNA substrate bound by PolY was plotted and fit to a sigmoidal curve by **Equation 2**.
